# A neural circuit algorithm for action grammar

**DOI:** 10.64898/2026.10.06.757194

**Authors:** Yue Liu, Lucas Y. Tian, Joshua B. Tenenbaum, Winrich A. Freiwald, Xiao-Jing Wang

## Abstract

How does the brain generate sequences of discrete elements, such as actions or words, that follow an abstract rule (a grammar)? Motivated by an experiment in which monkeys learned to draw multi-object images according to grammatical rules, we trained a recurrent neural network to generate sequences *A*^*m*^*B*^*n*^: producing shape A *m* times, then B *n* times, or vice versa. Our findings are threefold. First, the network plans the sequence by moving to an initial state encoding *m* and *n* along orthogonal directions for upcoming execution. Second, the network repeats each shape by progressing along a shape-specific manifold. Third, transitions from *A*^*m*^ to *B*^*n*^ follow parallel trajectories between manifolds across values of *n*, thus preserving *n*. Each of these is an experimentally testable prediction. The network generalizes to longer novel sequences by starting further along the same orthogonal directions. These results reveal a neural circuit implementation of a symbolic algorithm for generating rule-based action sequences.

## 1 Introduction

A hallmark of intelligence is the ability to generate unlimited new behaviors from a finite set of abstract rules, or grammars. Grammars specify how discrete elements, such as actions or words, combine into sequences (Fodor and Pylyshyn, 1988; Marcus, 2003; Dehaene et al., 2015, 2022), and are a fundamental building block of classic cognitive models of intelligent problem-solving (Newell, 1994; Anderson et al., 2004; Laird, 2012). Despite great progress in modern artificial intelligence (Goodfellow et al., 2016; Vaswani et al., 2017; Brown et al., 2020; Achiam et al., 2023; Singh et al., 2026; Guo et al., 2025), humans and animals remain more efficient than these systems at flexibly applying learned rules to new problems (Lake et al., 2015, 2017; Srivastava et al., 2023; Chollet et al., 2025; ARC Prize Foundation, 2026). The frontal and parietal cortices are implicated in organizing cognitive steps into a mental program using abstract grammatical rules (Duncan, 2010; El-Gaby et al., 2024; Tian et al., 2024; Manakov et al., 2026). However, how neural circuits implement grammatical rules to solve new problems remains poorly understood.

This work presents a neural circuit mechanism for grammar-based behavior in action sequencing. In particular, we focus on drawing behavior, which provides a useful platform to study how a sequence of actions is guided by structured knowledge (Lake et al., 2015; Tian et al., 2020; Fan et al., 2023). In a recent drawing task (Tian et al., 2026b), subjects (macaque monkeys) need to copy an image that is composed of multiple objects, and to do so by producing a sequence of strokes (one for each object) consistent with a particular set of sequencing rules (or action grammar). For example, during a particular trial, the subject may be presented with an image that contains 2 lines and 1 circle, and the action grammar could be “lines → circles”. In this case, the subject needs to first trace the 2 lines, followed by tracing the circle in order to obtain reward (Figure 1a). Therefore, the action sequences are always in the form of “*A*^*m*^*B*^*n*^”, meaning “producing shape A *m* times, and then shape B *n* times”.

**Figure 1:**
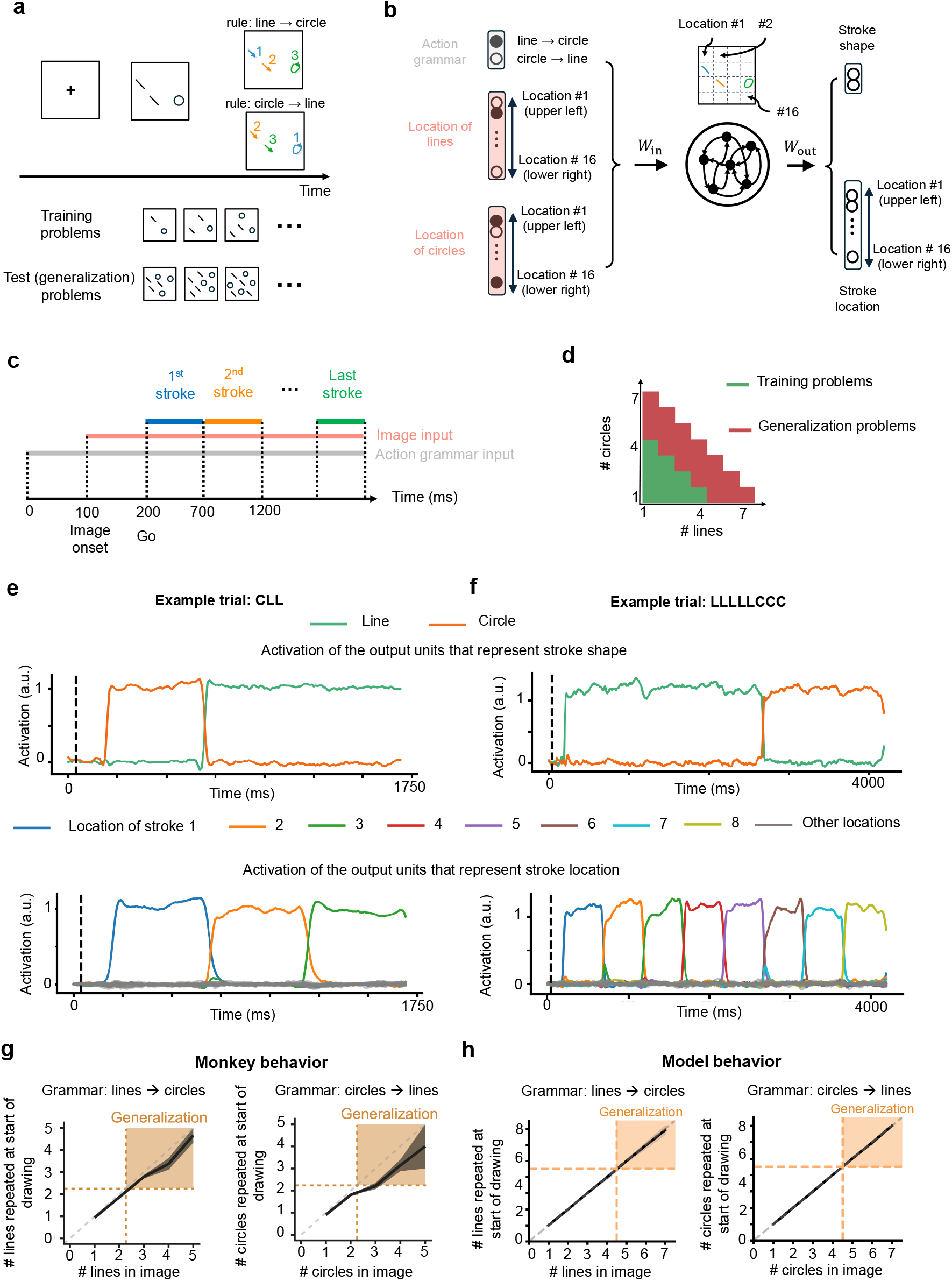
Trained recurrent neural network generalizes the action grammar. **a**. The schematic of the behavioral task in Tian et al. (2026b). Monkeys are presented with an image that contains multiple objects (lines and circles) and need to trace them using an abstract rule (for example, first trace all the lines and then all the circles). This rule is referred to as “action grammar”. Over training, each monkey also developed a spatial bias, for example tracing from top to bottom and left to right. Monkeys were trained using problems with a smaller number of objects and tested on problems with more objects, to see if they can generalize the learned action grammar to more difficult problems. **b**. A recurrent neural network (RNN) was trained to perform a simplified version of the monkey task. The RNN receives binary inputs about the grammar (“line → circle” or “circle → line”) and the locations of the lines and circles on the screen. The screen was discretized into a 4 × 4 grid to represent the locations. The RNN was trained to output a sequence that at each moment represents the shape and location of the current stroke. Each stroke lasts 500 ms. The input projects to all RNN units via the input weight (*W*_in_) and the outputs are read out from all RNN units via the output weight (*W*_out_). **c**. The temporal structure of the task used to train the RNN. **d**. The RNN was trained on problems that contain 2-5 objects. After that, all weights were frozen and the RNN was tested on problems that contain 6-8 objects. All problems contain both shapes. **e**. The activity of the RNN output units (top: shape, bottom: location) for an example training trial where it needs to produce a circle followed by two lines. **f**. The same as **e** but for a generalization trial where the RNN produced five lines followed by three circles. **g-h**. Performance of monkey and RNN across all training and generalization problems. Monkey data is reproduced from Tian et al. (2026b).

Importantly, subjects were not explicitly trained to use the action grammar. Nevertheless, they induced the action grammar from training. This is evident in a separate set of generalization trials, where they were presented with new problems that contain a larger number of objects. It was found that they can perform these novel, more complex problems by following the action grammar that they induced from training (Tian et al., 2026b). This behavioral result suggests that primates may have an inductive bias towards learning and using action grammars. Therefore, understanding the mechanisms by which neural circuits implement action grammars may generate broader insight into the neural basis of grammar-based cognitive behavior.

To explore potential neural circuit mechanisms behind this ability, we trained a recurrent neural network (RNN) on this task. We found that the trained RNN can generalize to novel, more complex problems, similar to monkeys. By reverse-engineering the trained RNN, we found three ingredients that together form an algorithm for solving the action grammar task. First, the state of the RNN before drawing encodes the number of objects for each shape in the image along orthogonal directions; second, the RNN state moves along these orthogonal directions during drawing different shapes. Finally, during the transition from drawing one shape to the next, the RNN state travels along parallel trajectories across trials, so that information about the number of repetitions for the upcoming shape is preserved. Together, these three ingredients form a symbolic algorithm consisting of two successive loops, where the first loop implements repetition of an action while the network maintains the information used by the second loop. This neural algorithm enables the RNN to generalize to novel problems by simply starting at an appropriate initial state according to the number of repetitions for each shape and the action grammar.

## 2 Results

### 2.1 Trained recurrent neural network generalizes the action grammar

We trained a recurrent neural network (RNN) to perform the action grammar task. The RNN receives inputs about the shape and location of all the objects on the screen, as well as an input about the rule for the current trial (Figure 1b, c). Each problem contains two shapes: lines and circles, and the RNN performs one of two rules during each trial: first produce lines then circles (“lines → circles”), or first produce circles then lines (“circles → lines”). The RNN was trained to output a sequence that represents the location and shape of each stroke under the rule (Figure 1b, c). Within the same shape, the RNN needs to produce the strokes from top to bottom, left to right, consistent with the observation that monkeys developed directional biases during training (Tian et al., 2026b). The inputs and target outputs are represented as binary vectors (Figure 1b, Methods).

The RNN was trained using supervised learning and backpropagation through time, on problems that contain 2-5 objects. After the RNN reached good performance on the training problems, we examined whether the RNN can generalize to more complex problems that contain 6-8 objects (Figure 1d). The RNN reached good performance on both the training and generalization problems, similar to monkeys (Figure 1e-h). In the following sections we will dive into the RNN to understand the mechanism that it uses to generate action sequences governed by abstract rules.

### 2.2. Repetition of different shapes is associated with movement along orthogonal manifolds

To understand the mechanism that the RNN uses to perform this task, we started by visualizing the RNN activity in a low-dimensional space. The basis axes of this space were defined by using linear regression to identify two directions in the state space that best represent the number of remaining lines and circles (Figure 2a, “repeat line” and “repeat circle” axes, see Methods for details), and a third direction that best discriminates the neural activity during drawing different shapes (Figure 2a, shape axis). These three axes were then orthogonalized (Methods).

**Figure 2:**
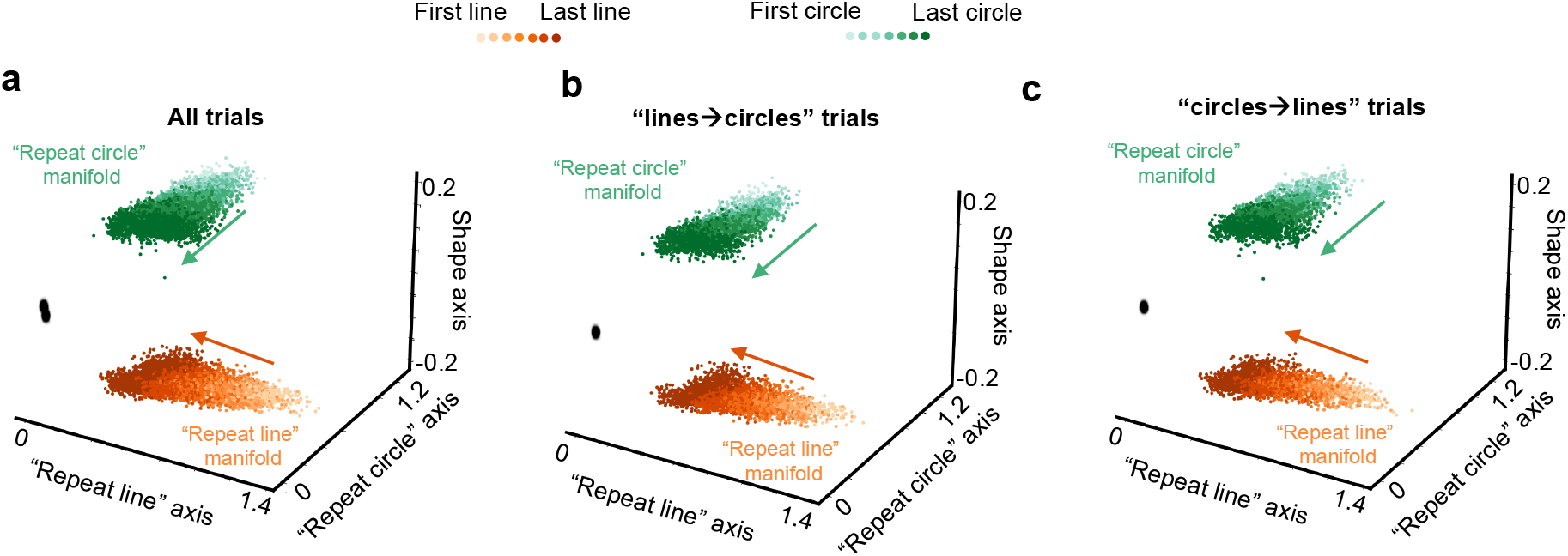
Repetition of different shapes is associated with movement along orthogonal manifolds. **a**. The RNN activity across all trials plotted in the low-dimensional space. Each point represents the average RNN activity during one stroke. Color represents the shape of the stroke. Darkness of the color represents the ordinal rank of the stroke within the repetition of the same shape. Darker color represents later strokes. Arrows indicate the direction that the neural state moves during repetition of lines and circles. **b**. The same result but for all “lines→circles” trials. **c**. The same result but for all “circles→lines” trials.

When projecting the time-averaged RNN activity during each stroke across all trials down to this low-dimensional space, two features stand out (Figure 2a). First, the RNN states occupy distinct parts of the state space while drawing different shapes. Second, as the RNN repeats a given shape, its state gradually moves along the corresponding repeat axes. Therefore, we defined the set of time-averaged RNN states while producing lines and circles “repeat line” and “repeat circle” manifolds. These repeat manifolds are reused across action grammars (Figure 2b-c). During each trial, the RNN state traverses these two manifolds sequentially in the order given by the action grammar. In the following sections, we will examine the detailed structures in the population dynamics while the RNN state traverses these two manifolds to reveal a network mechanism for this task.

### 2.3 Intuition for the network mechanism from three example trials

To perform this task, the RNN needs to transition from one repeat manifold to the other upon finishing the first shape. In order to make this transition at the correct time, the RNN needs to keep track of the remaining number of actions until the transition. To build intuition for how this is achieved, we first examined neural trajectories during three example trials where the network performs the “circles → lines” rule with different numbers of repetitions for lines and circles (Figure 3).

**Figure 3:**
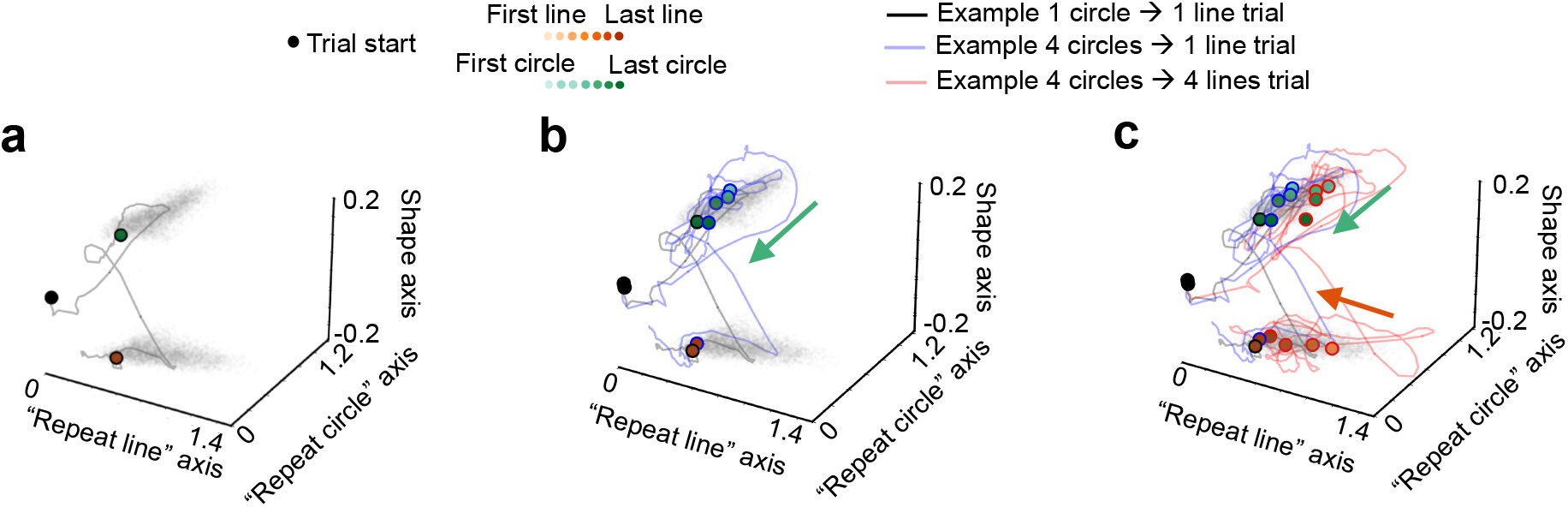
Neural trajectories during three representative trials. The neural trajectories during example trials where the RNN produces 1 circle followed by 1 line (“1 circle→ 1 line” trial, gray lines), 4 circles followed by 1 line (“4 circles→ 1 line” trial, blue lines), and 4 circles followed by 4 lines (“4 circles → 4 lines” trial, red line). Comparing the “1 circle 1 line” and “4 circles→ 1 line” trials (**b**), the initial state for the “4 circles→ 1 line” trial (the lightest green circle on the blue trajectory) is shifted along the “repeat circle” axis, such that it travels a longer distance during circle repetitions. Comparing the “4 circles→ 1 line” and “4 circles→ 4 lines” trials (**c**), the initial state of the “4 circles→ 4 lines” trial (the lightest green circle on the red trajectory) is further shifted along the “repeat line” axis, such that it will travel a longer distance during line repetitions after transitioning to the “repeat line” manifold. As a reference, the “repeat line” and “repeat circle” manifolds in Figure 2 are plotted using gray circles in the background.

Figure 3a shows the neural trajectory during a trial where the RNN needs to produce 1 circle followed by 1 line. After the trial starts, the RNN state travels to the edge of the “repeat circle” manifold and produces a circle, before making a transition to the edge of the “repeat line” manifold and producing a line.

Figure 3b shows the same trial as in Figure 3a. In addition, it also shows the neural trajectory during another trial where the RNN produces 4 circles followed by 1 line. Compared to the “1 circle → 1 line” trial, the RNN state starts further along the “repeat circle” axis when producing the first circle during the “4 circles→ 1 line” trial (compare the lightest green circle along the blue trajectory to the green circle along the gray trajectory). During the repetition of circles, it travels along the “repeat circle” axis (towards smaller values). The rest of the dynamics are similar to the “1 circle→ 1 line” trial: as the RNN state reaches the edge of the “repeat circle” manifold, it transitions to the “repeat line” manifold, before landing on the edge of the “repeat line” manifold and producing one line.

Figure 3c shows the neural trajectory during an additional trial where the RNN produces 4 circles followed by 4 lines. Compared with the “4 circles→ 1 line” trial, the RNN state starts at about the same location along the “repeat circle” axis, but further along the (orthogonal) “repeat line” axis (compare the lightest green circle along the red trajectory to those along the gray and blue trajectory). It then travels along the “repeat circle” axis (towards smaller values) while the RNN produces 4 circles. As it reaches the edge of the “repeat circle” manifold, the location of the RNN state remains further along the “repeat line” axis compared with the “4 circles→ 1 line” and “1 circle→ 1 line” trials (compare the darkest green circle along the red trajectory to those along the gray and blue trajectory). Then, the RNN state transitions to the “repeat line” manifold. After the transition, the state is still further along the “repeat line” direction. It then travels along the “repeat line” direction (towards smaller values) to produce 4 lines.

A few observations can be made from these example trials which will be tested more rigorously in later sections. First, the RNN state transitions to the next shape when it reaches the edge of the repeat manifold for the first shape, regardless of the number of repetitions it has completed. Second, the RNN state starts further away from the edge of the first repeat manifold if it were to produce more repetitions for the first shape. This allows the RNN to travel a longer distance before reaching the edge of the manifold where it will transition to the next shape. Third, the number of repetitions of the second shape is also represented when the network draws the first shape, but along an orthogonal direction. As a result, if there are more repetitions for the second shape, the RNN state will remain further along the repeat axis for the second shape during drawing the first shape. Finally, the transition to the second repeat manifold preserves this representation of the second shape, so that if the RNN state is further along the repeat axis for the second shape before the transition, it will be further away from the edge of the second repeat manifold after the transition. Consequently, the RNN can travel a longer distance along the second repeat manifold in order to produce more repetitions for the second shape.

In the following two sections we will test the conclusions from the intuitive picture above. First, we will test whether the initial state of the RNN represents the number of lines and circles in the image along orthogonal directions. Second, we will examine if the transition between the two repeat manifolds preserves the number of repetitions for the upcoming shape.

### 2.4 Orthogonal neural dimensions encode the number of each shape before drawing

Our first observation from examining the RNN dynamics during example trials (Figure 3) is that the initial state of the RNN before drawing represents the number of lines and circles along orthogonal directions. To test this more rigorously, we plotted the RNN state at the Go cue across trials in the low-dimensional space spanned by the “repeat line” and “repeat circle” axes. We indeed found that the initial state represents the number of lines and circles along the corresponding repeat axes (Figure 4a-b). Importantly, this organization also extends to the generalization trials, where the RNN produces more repetitions than during training (crosses in Figure 4a).

**Figure 4:**
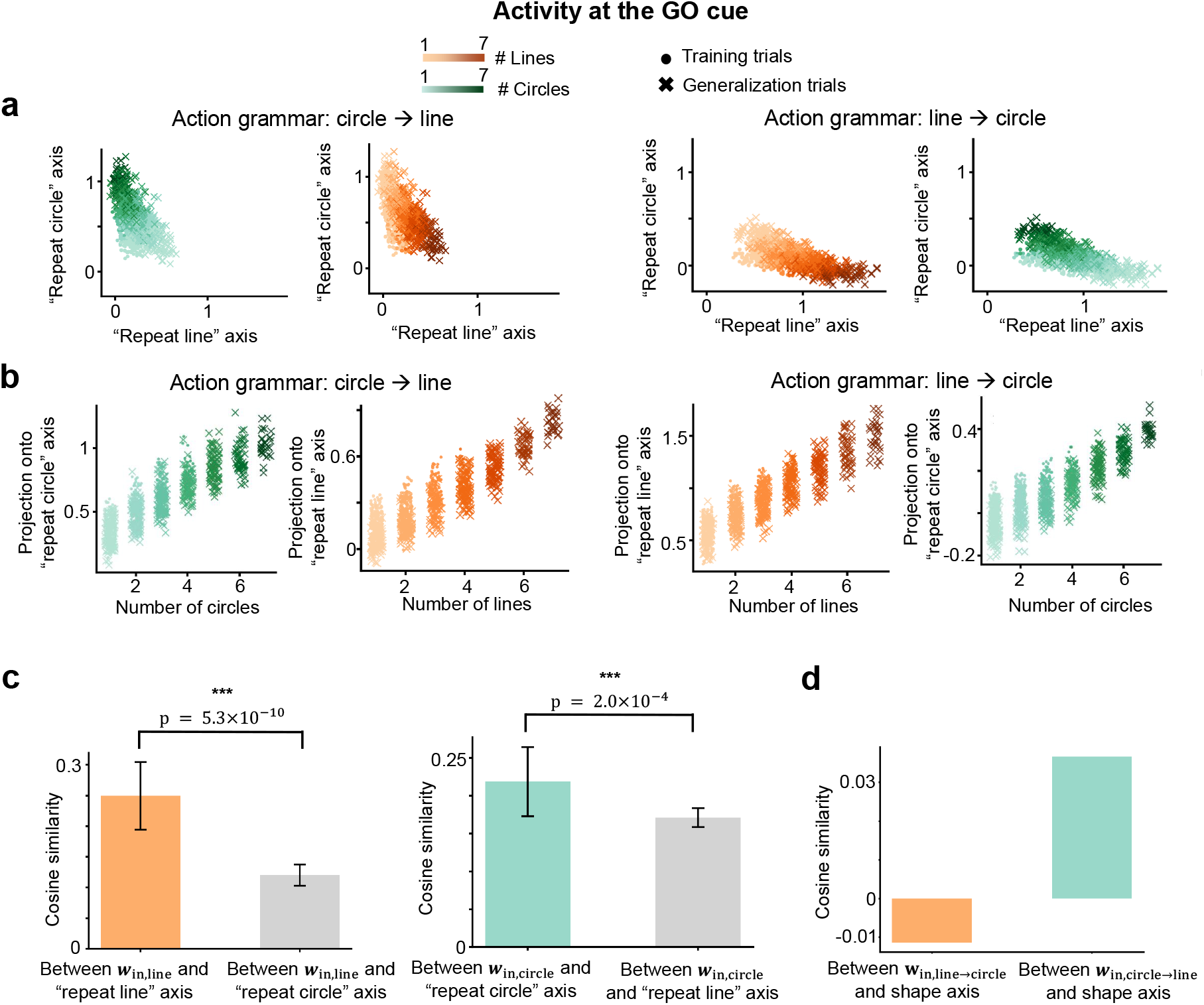
Orthogonal neural dimensions encode the number of each shape before drawing. a. The activity of the network at the Go cue, plotted in the two-dimensional space spanned by the “repeat line” and “repeat circle” axes. Each point represents a trial and is colored in one of two ways: according to the number of lines (orange gradient) or the number of circles (green gradient) in the image. Circles represent training trials and crosses represent generalization trials. It can be seen that the activity states before drawing represent the number of lines and circles along the orthogonal “repeat line” and “repeat circle” directions, and this relationship also holds for the generalization trials. b. The projection of the network state at the Go cue onto the “repeat line” and “repeat circle” axes, for trials with different numbers of lines and circles. A small noise was added to the plotted *x* values for visual clarity. c. The cosine similarity between the input weight vectors originating from input units that represent the locations of lines (left) / circles (right) and the “repeat line” and “repeat circle” axes. The input weight vectors originating from input units representing lines are preferentially aligned with the “repeat line” axis (one-sided t-test, *p* = 5.3 × 10^*−*10^); similarly, input weight vectors from units representing circles are more aligned with the “repeat circle” axis (one-sided t-test, *p* = 2.0 × 10^*−*4^). This structure in the input weights leads the initial network state to the appropriate location on the repeat manifold. d. The weight from the input unit that represents the lines → circles action grammar is anti-aligned with the shape axis (the vertical axis in Figure 2a); conversely, the weight from the input unit that represents the circles → lines grammar is aligned with the shape axis. This structure in the input weights pushes the initial network state to different repeat manifolds depending on the action grammar for the current trial.

We next sought to answer what structural properties of the RNN enable the initial state to represent the number of lines and circles along orthogonal dimensions. Because the time between trial onset and the start of movement is quite short (Figure 1c), we hypothesized that the external input plays an important role in moving the RNN state to the appropriate location in the state space before drawing starts. Indeed, when examining the input weights, we found that the connection weight vectors from input units that represent the location of lines (circles) are more aligned with the “repeat line” (“repeat circle”) axes compared to the “repeat circle” (“repeat line”) axis (Figure 4c). Therefore, if an image contains more lines (circles), the RNN will receive a stronger input along the “repeat line” (“repeat circle”) direction. As a result, the network state will be more strongly pushed along the corresponding repeat axis before drawing starts.

Moreover, the weight vector from the input unit that represents the “circles → lines” rule is aligned with the shape axis, whereas the weight vector from the “lines → circles” input unit is anti-aligned with the shape axis (Figure 4d). This property of the input weight ensures that the RNN state is pushed upwards in the low-dimensional space (see Figure 2 and 3) when the rule is “circles→ lines”, and downwards when the rule is “lines→ circles”.

In summary, the structures in the input weights enable the RNN state to start at the appropriate location on the correct repeat manifold. This initial state represents a high-level plan for the upcoming execution of the sequence.

### 2.5 Parallel dynamics during shape transition preserve the number of repetitions for the upcoming shape

From the example trials in Figure 3c, we see that the RNN state maintains the number of the second shape along the corresponding repeat axis while drawing the first shape. Importantly, this representation is also observed immediately after transitioning to the second shape, which enables the RNN to produce the correct number of the second shape. How does the RNN preserve the information about the number of the upcoming shape during the transition between shapes?

One way to preserve the information is for the neural trajectories during the transition to be parallel across different trials, so that the orderly organization of the states before the transition is preserved after the transition. To test this hypothesis, we first visualized the neural trajectories during the transition, colored by the number of repetitions of the upcoming shape (Figure 5a-b). From visual inspection they indeed seem close to parallel. To test this more rigorously, we computed the average cosine similarity between the tangent vectors along pairs of trajectories, which represents the degree of parallelism between them at each point in time. The average cosine similarity is significantly higher than chance throughout the transition (Figure 5c-d). This shows that the RNN uses parallel dynamics to preserve the information about the number of repetitions for the upcoming shape during the transition between shapes.

**Figure 5:**
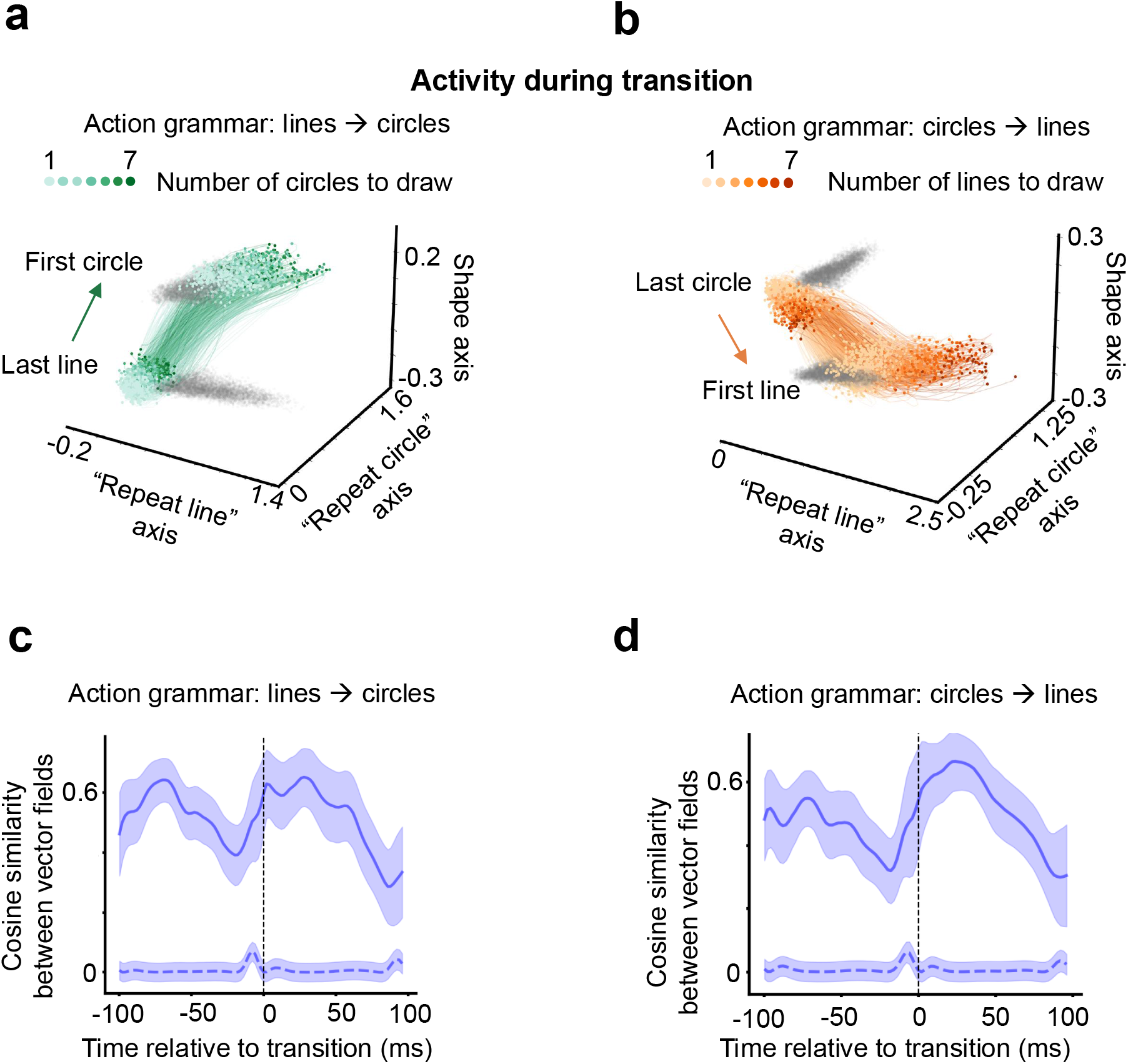
Parallel dynamics during shape transition preserve the number of repetitions for the upcoming shape. a. The network dynamics during the transition from producing lines to producing circles (50 ms before finishing last line to 50 ms after starting the first circle). Color represents the number of circles that the network is going to draw after the transition. The ordered representation about the number of upcoming circles is preserved during the transition. b. Same as **a** but for the transition from the last circle to the first line. c. Cosine similarity between velocity vectors along pairs of neural trajectories that represent different trials, during the transition from lines to circles. Dashed line: the same cosine similarity computed when the identity of the RNN units was randomized. Shaded region represents standard deviation across trials. Neural trajectories during the transition are significantly more parallel than chance, which preserves the orderly organization of neural states before the transition. d. The same as **c** but for the transition from circles to lines.

### 2.6 Single neurons encoding shape, ordinal rank, and sequence boundary

Finally, we turned to look at the responses of single RNN units. Based on the population-level picture above, we hypothesized that there should be at least three types of single unit responses in the RNN. First, some units should encode the shape of the current stroke that the network is producing. This response creates the separation between the two repeat manifolds along the shape axis in Figure 2a. Indeed we found many units whose activity is strongly modulated by the current shape (Figure 6a). We further investigated the dynamics of the shape-modulated units around the time of shape transition. During this time, the activity of shape-selective units that prefer the first shape rapidly decreases, and that of shape-selective units that prefer the second shape rapidly increases (Figure 6b). What could trigger this rapid transition? We found that the inhibitory input from units selective for the first shape to units selective for the second shape progressively weakens over the strokes leading up to the transition (Figure 6c). This gradual release from inhibition may allow the second-shape-selective units to become active once the first shape is complete.

**Figure 6:**
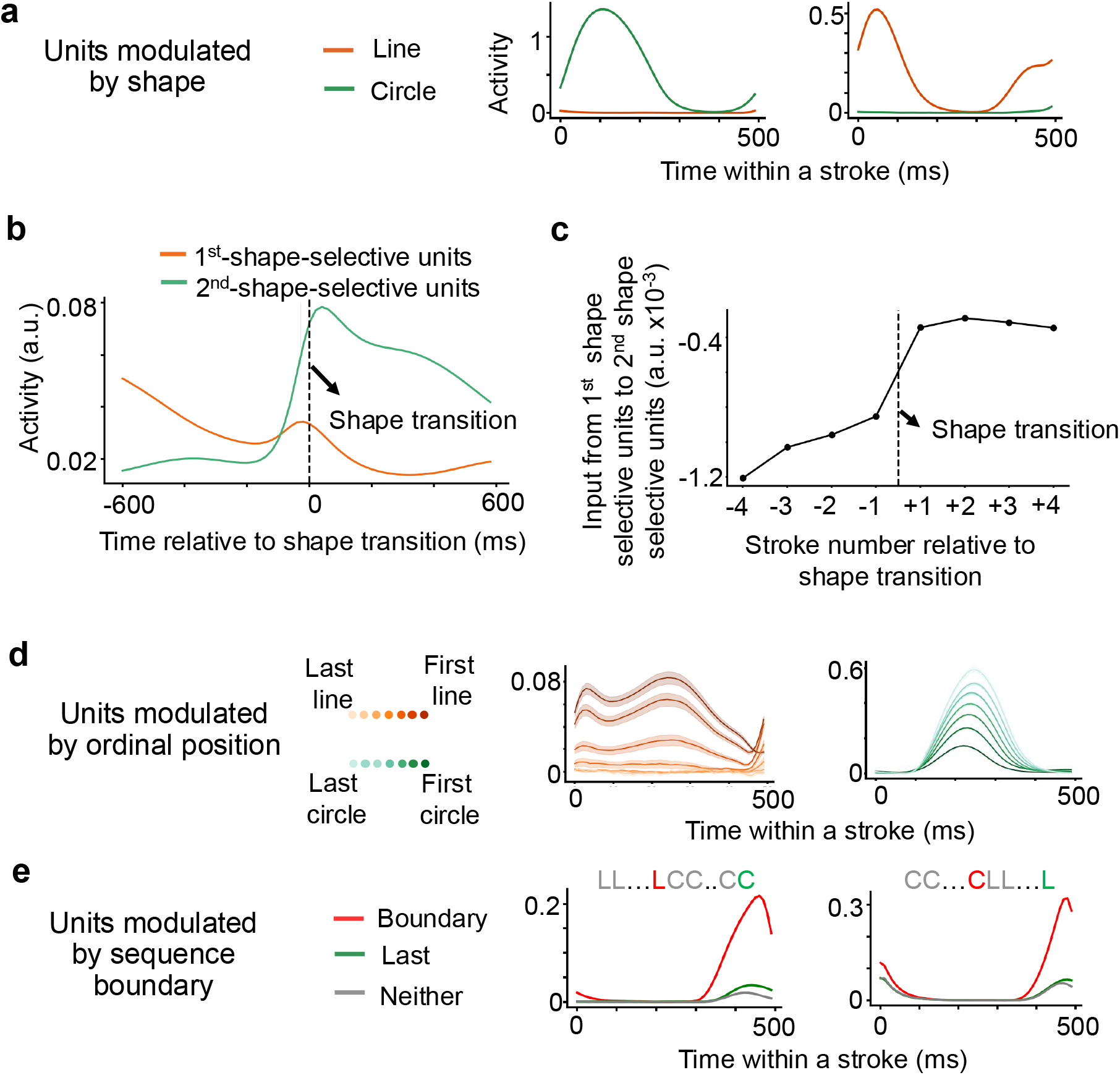
Single units in the RNN encoding shape, ordinal rank, and sequence boundary. a. Example single units in the RNN that have differential activity depending on the shape of the current drawing. b. Average activity of RNN units that are selective to the first and second shape around the transition between shapes. The activity of RNN units selective for the second shape sharply increased as the network switches to drawing that shape. c. Mean input current from RNN units selective for the first shape to those selective for the second shape as a function of stroke number relative to the transition. The inhibition from the first-shape-selective units to the second-shape-selective units was reduced as the transition time was approaching. d. Example single units in the RNN that have differential activity depending on the ordinal position of the current drawing. e. Example single units in the RNN that are especially active before the transition to drawing the next shape. The red curve represents the activity during the last stroke before the transition. The green curves represent the activity during the very last stroke of the sequence. The gray curves represent the activity during other strokes. L: line; C: circle. For **a, d, e**: Solid lines represent trial-averaged activity. Shaded areas represent standard error of the mean.

Besides shape-modulated units, the RNN also contains many units that are modulated by the current stroke’s ordinal position within the repetition (Figure 6d). These units underlie the progression along the shape-specific repeat manifolds (see Figure 2a).

Finally, there are also units that are especially active at the “action sequence boundary”, just before the transition to the next shape (Figure 6e). Their timing suggests that they help drive the network state between the two repeat manifolds during the transition (Figure 5).

## 3 Discussion

In this work we studied how neural networks can generate sequences that follow abstract rules of the form *A*^*m*^*B*^*n*^, where *A* and *B* are distinct actions, and *m* and *n* are the number of times each action should be repeated. We found that the dynamics of the trained RNN implement an algorithm, which is summarized in Figure 7 using an example “circles→lines” trial.

**Figure 7:**
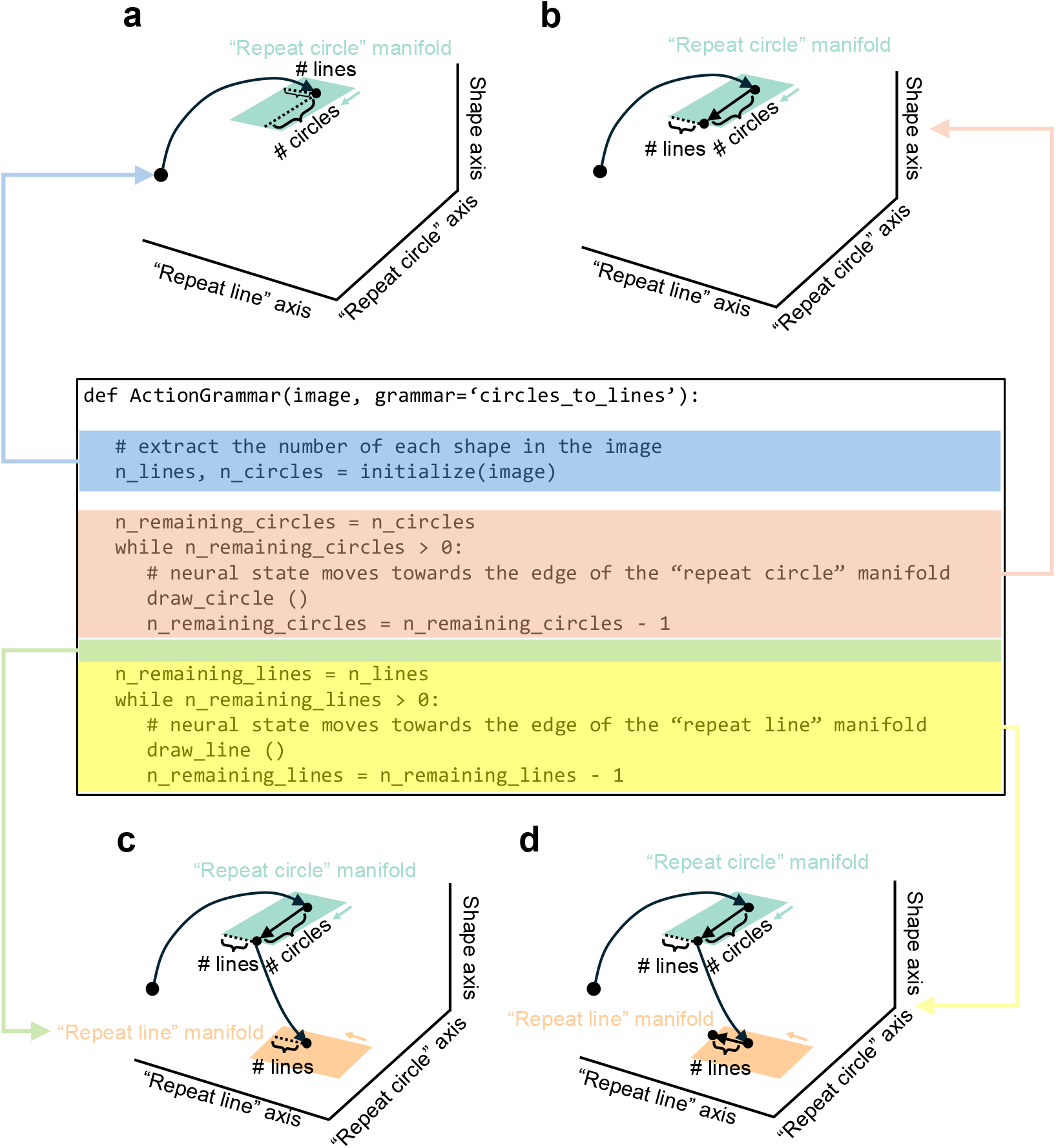
A schematic of the discovered neural algorithm for generating sequences of the form *A*^*m*^*B*^*n*^. The mechanism used by the RNN to perform the task, illustrated using an example circles → lines trial. a. When presented with the image, the neural state moves to a particular location on the “repeat circle” manifold, where the distances to the two edges of the manifold are proportional to the number of lines and circles in the image. b. The neural state gradually moves along the “repeat circle” axis to the edge of the “repeat circle” manifold to produce the correct number of circles. At this point, the distance to the other edge of the manifold represents the number of lines left to draw. c. The neural state transitions to the “repeat line” manifold. Importantly, the distance to the edge of the manifold is preserved via parallel dynamics (see Figure 5), such that it still represents the number of lines to draw along the “repeat line” axis after the transition. d. The neural state then gradually moves towards the edge of the “repeat line” mani-fold to produce the correct number of lines.

- At the beginning of the trial, the external input drives the RNN state to a specific location on the “repeat circle” manifold. The distances to the two edges of the manifold represent the number of lines and circles to draw, respectively (Figure 7a).
- The RNN produces the correct number of circles by gradually moving along the “repeat circle” direction on the “repeat circle” manifold, until it reaches the edge of the manifold. At this time, the distance to the other edge of the manifold represents the number of lines to draw next (Figure 7b).
- The RNN transitions from the “repeat circle” manifold to the “repeat line” manifold. This transition is parallel across different trials, therefore it preserves the information about the number of lines. As a result, after the transition, the distance to the edge of the “repeat line” manifold also represents the number of lines to draw next (Figure 7c).
- The RNN produces the correct number of lines by gradually moving along the “repeat line” axis until it reaches the edge of the “repeat line” manifold (Figure 7d).

In principle, this mechanism can generate any action sequence in the form of *A*^*m*^*B*^*n*^ by setting the initial network state at the appropriate location on the repeat manifold^1^. In other words, it implements an algorithm for generating action sequences in the form of *A*^*m*^*B*^*n*^. Therefore, this work bridges the gap between symbolic algorithms and their implementation in neural networks—a longstanding question in cognitive science, neuroscience, and artificial intelligence (Lake et al., 2017; Marcus, 2003; Smolensky, 1990). This work also provides a concrete mechanism for how neural circuits organize a series of processing steps to implement a mental program (Duncan, 2010; Tian et al., 2024; Dehaene et al., 2022).

An important part of the mechanism is that the initial state encodes the number of repetitions for each shape in the action sequence along orthogonal directions. This is reminiscent of the neural activity patterns that predict multiple future movements observed in the frontal cortex of macaque monkeys (Averbeck et al., 2002; Mushiake et al., 2006; Shima et al., 2007; Xie et al., 2022; Tian et al., 2024; Krasa et al., 2026). In particular, in the study of Shima et al. (2007), it was shown that this activity represents abstract structural properties of the future action sequence (whether it is of the form “AABB”, “ABAB”, or “AAAA”). However, it is not well understood how this abstract action plan contributes to the subsequent generation of action sequences which follow abstract rules. The present work provides such a mechanism, demonstrating how this pre-movement activity pattern combines with subsequent dynamics to implement the action grammar.

Our results provide insight into the relationship between single unit activity and population dynamics. We observed single units in the RNN that encode the shape and ordinal rank of the current stroke. In addition, some units are particularly active before the action sequence boundary when the network is about to transition to the next shape (Figure 6). Similar single neurons have been discovered in the frontal cortex of monkeys performing motor sequencing tasks. For example, single neurons that encode the shape of the current drawing in an abstract format (invariant to size and location) have recently been discovered in the ventral premotor cortex when monkeys perform a similar drawing task (Tian et al., 2026a). Single neurons that encode the ordinal rank of an action in a sequence have been discovered in prefrontal and premotor cortices (Clower and Alexander, 1998; Shima and Tanji, 2000; Mushiake et al., 2006; Berdyyeva and Olson, 2010) and implemented in neural network models for action sequencing (Salinas, 2009). Neurons that represent action sequence boundaries have also been reported in the prefrontal cortex (Fujii and Graybiel, 2003). Therefore, our mechanism provides an account for how these different populations of neurons in the frontal lobe might work together to generate action sequences that satisfy abstract rules.

Alternative mechanisms for generating action sequences in the form of *A*^*m*^*B*^*n*^ are possible. For example, while our model encodes the number of repetitions as the distance to the edge of the repeat manifold, it could also be implemented by the *speed* at which the neural state moves towards the edge of the repeat manifold, such that the network state moves at a slower speed if it needs to repeat more actions. However, such a speed-based mechanism would likely be more difficult to realize in neural networks, as it would require rescaling the neural dynamics whenever the network faces a new problem. In contrast, the distance-based mechanism we observed allows dynamics to be reused across problems, which potentially facilitates more efficient generalization.

In addition, our RNN plans out the entire future action sequence by mapping the sequence to an appropriate initial network state using input-driven dynamics (Figure 4). While this mechanism is straightforward, other planning mechanisms may also be possible, such as forward replay (Johnson and Redish, 2007; Pfeiffer and Foster, 2013; Singer et al., 2013) and recurrent dynamics (Jensen et al., 2024).

The RNN used in this study receives abstract visual inputs about the shape and location of each object on the screen, and generates abstract motor commands representing the shape and location of each stroke in the action sequence. In other words, it focuses on the transformation from high-level visual inputs to abstract motor commands. As such, it does not address the transformation at the sensory and motor ends, namely how the pixel-level information about the image is transformed into the abstract representation about the shape and location of each object in the image, and how the abstract representation of an action (about drawing a particular shape at a particular location) is transformed into moment-by-moment movement kinematics. Incorporating more realistic visual front end and motor back end is an interesting direction for future research.

Generating sequences of the form *A*^*m*^*B*^*n*^ is only one simple case of grammar-based behavior. Natural grammars, such as those of language, involve richer structures, including nested and hierarchical dependencies (Dehaene et al., 2015). Future work could investigate how neural circuits implement more complex grammars, and how they assemble learned primitives, such as the repetition loop studied here, into new mental programs (Lake et al., 2017; Lake and Baroni, 2023; Tian et al., 2020; Lake et al., 2015; Rule et al., 2020; Ellis et al., 2023). A mechanistic understanding of how neural circuits build and execute mental programs is critical for understanding the flexibility of cognition, and could also inspire the development of more adaptive and data-efficient artificial intelligence systems.

## 4. Methods

### 4.1 The RNN model

We used a vanilla RNN with *N* = 1024 units. The network activity evolves according to the equation

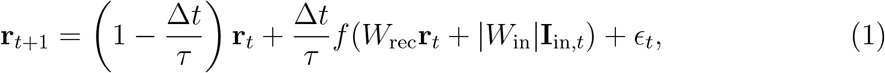

where vector **r**_*t*_ is the activity of all units in the RNN at time step *t*, Δ*t* = 25 ms is the duration for each timestep, *τ* = 100 ms is the decay time constant of the neurons, *W*_rec_ is the recurrent weight matrix, *W*_in_ is the input weight matrix. We take the element-wise absolute value of the input weight matrix to reflect the fact that long-range connections in the brain are predominantly excitatory. **I**_in_ is the input current, and *ϵ* is the noise of the network sampled from a Gaussian distribution 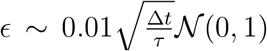 is the nonlinear input-output function, and is chosen to be a ReLU function.

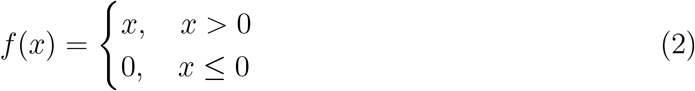

The RNN generates an output sequence through

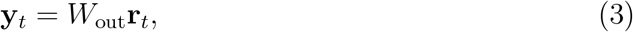

where *W*_out_ is the output weight matrix.

All trainable parameters were initialized using orthogonal matrices as described in Saxe et al. (2013).

### 4.2 Task

We trained the RNN to perform an abstract version of the action grammar task in Tian et al. (2026b). During each trial, the RNN receives an input about the shape and location of each object on the screen. The screen is discretized into a 4 × 4 grid so that each object is assigned to one of sixteen locations. At each time point, this input is represented as a 32-dimensional vector. The first 16 entries represent the location of the lines, and the next 16 entries represent the location of the circles. An entry is 1 if there is a line or circle at the corresponding location, and 0 otherwise. This input **I**_image_(*t*) is constant from 100 ms after trial onset to the end of the trial. Mathematically, for *k* ∈ {1, 2, 3, …, 16},

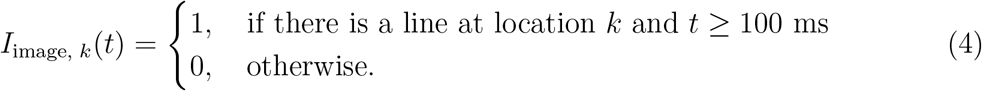

For *k* ∈ {17, 18, 19, …, 32},

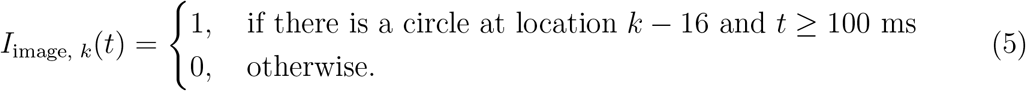

The RNN also receives input about the action grammar it needs to apply for the current trial. The action grammar is represented by a two-dimensional one-hot vector, where each entry corresponds to one of the two rules (“lines→circles” or “circles→lines”).

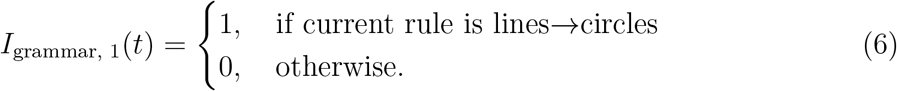

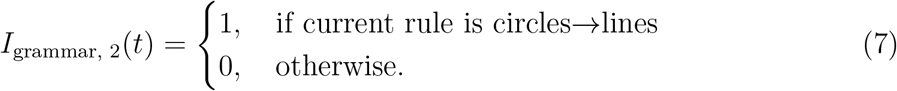

The grammar input is present throughout the trial.

The RNN also receives a brief go cue at *t* = 200 ms that lasts for 50 ms.

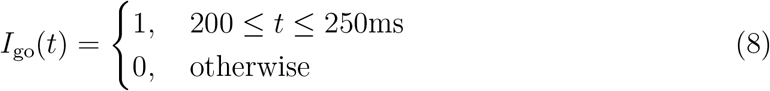

The total input to the RNN is the combination of the image, grammar, and go inputs.

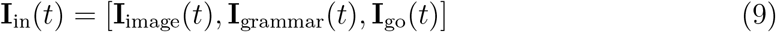

The target output of the RNN represents a sequence of strokes. At each time, it is an 18-dimensional vector. The first 16 dimensions represent the location of the current stroke, and the last 2 dimensions represent the shape of the current stroke. Each stroke lasts 500 ms.

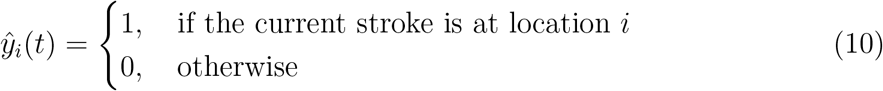

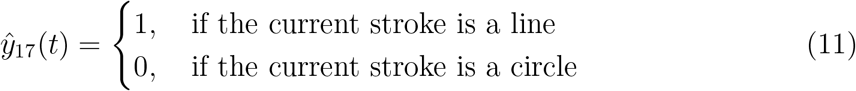

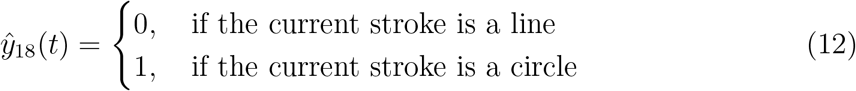

### 4.3 Training and generalization

We generated problems with 2-5 objects as the training set (each problem contains at least one object for each shape). The RNN was trained with supervised learning using the Adam optimizer (Kingma and Ba, 2014) with a learning rate of 0.001 and the mean-squared loss.

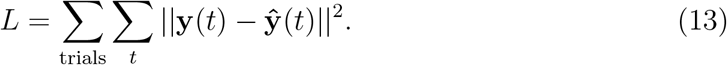

After training converged, we tested the RNN using problems that contain 6-8 objects.

### 4.4 Dimensionality reduction

Here we describe how the “repeat line”, “repeat circle” and the shape axes were obtained for dimensionality reduction. To obtain the repeat axes, we first computed the average activity of each RNN unit during each stroke. Then, linear regression models were built to predict the number of remaining strokes with the same shape (*n*_remain_) from this stroke-averaged activity. To isolate the encoding of *n*_remain_ from other task variables, we fitted a separate linear regression model for each unique combination of action grammar, shape, location, and ordinal rank^2^. Mathematically,

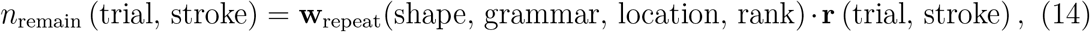

where **r** (trial, stroke) is the average population activity during a particular stroke within a particular trial, and · denotes the dot product. The vectors **w**_repeat_ turn out to mainly depend on shape, so we averaged them over all the other variables to obtain one repeat axis for each shape

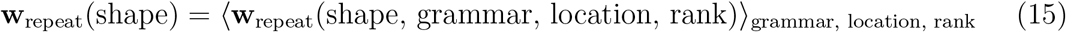

The shape axis was similarly obtained by fitting a linear model to predict the shape of each stroke using stroke-averaged network activity. The shape is represented as a binary variable

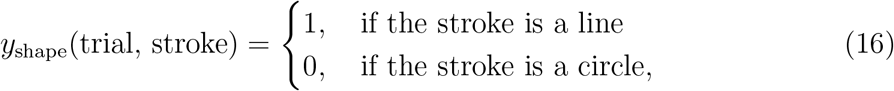

and the linear model is defined as

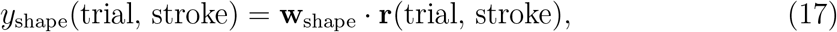

where **r**(trial, stroke) is the average population activity during a particular stroke within a particular trial.

All three axes were fit using stroke-averaged activity from all trials, including generalization trials. Finally, we used Gram-Schmidt orthogonalization on the set of vectors {**w**_repeat_(line), **w**_repeat_(circle), **w**_shape_} to obtain a set of orthogonal vectors used in the main figures.

### 4.5 Quantifying degree of parallelism between neural trajectories during the transition

We quantified the degree of parallelism between pairs of neural trajectories by computing the cosine similarity between the velocity vectors taken at the same time

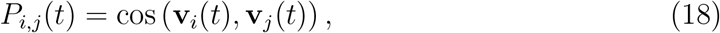

where *i* and *j* are indexed for the neural trajectories. Velocity was computed as the difference between the population vectors at consecutive time points

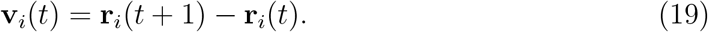

The average cosine similarity across pairs of neural trajectories was plotted in Figure 5c and d.

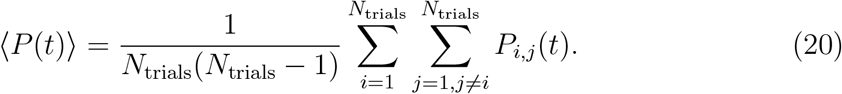

## Acknowledgements

We thank members of the Wang, Freiwald and Tenenbaum labs for discussions. This work was supported by the Office of Naval Research (N00014-20-1-2292, Vannevar Bush Faculty Fellowship, to W.A.F., N00014-23-1-2040 to X.-J.W., and MURI N00014-21-1-2801 to J.B.T.), the National Institutes of Health, through the National Eye Institute (R01EY021594 to W.A.F.), the National Institute of Mental Health (F32MH125573 to L.Y.T.), and the National Institute of Neurological Disorders and Stroke (K99NS131585 to L.Y.T.), as well as the Center for Brains, Minds and Machines of the National Science Foundation (STC award CCF-1231216 to W.A.F. and J.B.T.), and the Air Force Office of Scientific Research (FA9550-22-1-0387 to J.B.T.).

## Footnotes

1 In reality, the maximum *m* and *n* should depend on the length of the repeat manifold along the corresponding repeat direction

2 Notice that this is the “forward” rank (e.g. the first line, the second circle, etc.), which is different from *n*_remain_ that represents how many lines/circles are left to draw.

